# *Cfdp1* is Essential for Cardiac Development and Function

**DOI:** 10.1101/2022.03.21.485062

**Authors:** Giardoglou Panagiota, Deloukas Panos, Dedoussis George, Beis Dimitris

**Affiliations:** Zebrafish Disease Model Lab, Center for Clinical, Experimental Surgery & Translational Research, Biomedical Research Foundation, Academy of Athens, Athens, Greece; Department of Nutrition and Dietetics, School of Health Science and Education, Harokopio University of Athens, Athens, Greece; Clinical Pharmacology, William Harvey Research Institute, Barts and The London Medical School, Queen Mary University of London, London, UK

## Abstract

Cardiovascular diseases (CVDs) are the prevalent cause of mortality worldwide and account for the most common noncommunicable disease. CVDs describe a wide spectrum of disorders affecting the proper function, physiology and morphogenesis of the heart and blood vessels. The risk of developing cardiovascular diseases is modulated by a combination of environmental and genetic effectors. Thus, it’s highly important to identify candidate genes and elucidate their role in the manifestation of the disease. Large-scale human studies have revealed the implication of Craniofacial Development Protein 1 (*CFDP1)* in coronary artery disease (CAD). *CFDP1* belongs to the evolutionary conserved Bucentaur (BCNT) family and up to date, its function and mechanism of action in Cardiovascular Development is still unclear. In this study, we utilize zebrafish to investigate the role of *cfdp1* in the developing heart due to the high genomic homology, similarity in heart physiology and the ease of experimentally manipulation. We showed that *cfdp1* is expressed during development and at 120 hours post fertilization its expression is restricted to the region of the heart and the head. We then generated a *cfdp1*-null zebrafish line using CRISPR-Cas9 system which led to a lethal phenotype since *knockout* embryos do not reach adulthood. *cfdp1*^−/−^ embryos develop arrhythmic hearts and defective cardiac performance exhibiting statistically significant differences in heart features including End Diastolic Volume, Cardiac Output, Ejection Fraction and Stroke Volume. Myocardial trabeculation is also impaired in *cfdp1*^−/−^ embryonic hearts, implying its regulatory role also in this developmental process. Findings from both *knockdown* and *knockout* experiments showed that abrogation of *cfdp1* leads to downregulation of Wnt signaling in embryonic hearts during valve development but without affecting Notch activation in this process.

## INTRODUCTION

Cardiovascular diseases (CVD) comprise a broad spectrum of cardiac defects and clinical characteristics. Multiple factors contribute to the severity of CVD traits and it still remains unclear in which extend genetics together with environmental elements lead to the disease manifestation. The suggested predisposition of the CVD appearance is currently one the main focus of research interest. Genome-wide association studies have identified thousand robust associations (genome-wide significance, *p* < 5×10^−8^) between disease traits and genetic loci. To date, for coronary artery disease (CAD), 66 loci have been proposed to account for approximately 12% of CAD heritability^1–4^. Moreover, it has also been reported that a larger number of putative loci is found at the false discovery rate (FDR) of 5%. Recently, a study that used UK Biobank data^5^ to evaluate the validity of FDR loci and conducted meta-analysis using CAD GWAS identified new loci at GWAS significance that were previously on 5% FDR providing support that variants in this threshold could hold the key for higher percentage of heritability^4^.

Findings from human studies trying to analyze the genetic architecture of normal heart physiology, link the cardiac structure and function with SNPs (single nucleotide polymorphisms) in identified genetic loci. A list of recent publications derived from analysis of GWAS human data have highlighted the involvement of *CFDP1* (craniofacial development protein 1) in the determinants of risk factors for CAD, blood pressure, aortic diameter and carotid intima-media thickness raising the interest for deeper understanding of the functional analysis of *CFDP1* gene in cardiovascular development and function^6–9^.

The human *CFDP1* is a protein-coding gene belonging to the evolutionary conserved Bucentaur (BCNT) superfamily which is classified by the uncharacterized BCNT domain of 80 amino acids (aa) at the C-terminal region. It is located at the reverse strand of chromosome 16, consists of 7 exons, which encodes for a protein product of 299 aa and it is flanked by the *BCAR1* (breast cancer antiestrogen resistance 1) and *TMEM170A* (transmembrane protein 170A) genes. The BCNT protein family is widespread among the species and their orthologues are also found in yeast *Saccharomyces cerevisiae* (SWC5), fruit fly *Drosophila melanogaster* (YETI), mouse *Mus musculus* (CP27) and zebrafish *Danio rerio* (RLTPR or CFDP1)^10^. The fact that BCNT is evolutionary conserved implies the important role of this superfamily, which was first detected in bovine brain extracts using monoclonal antibodies against a rat GTPase-activating protein with the same epitope^11^. Although *CFDP1* gene is highly conserved, there is limited knowledge about its function and role not only in cellular level but also in level of organism. Yeast *Swc5* gene is essential for optimal function of chromatin remodeler SWR which has histone exchange activity in an ATP-dependent manner^12–14^. Studies in *Drosophila melanogaster* have shown that loss of BCNT gene, *Yeti* causes lethality before pupation and that mechanistically provides a chaperon-like activity that is required in higher-order chromatin organization by its interaction with histone variant H2A.V and chromatic remodeling machinery^15^. In the same context, functional analysis of zebrafish *cfdp1* has shown evidence for its role in craniofacial structure and bone development^16^. Mammalian BCNT proteins have also been characterized as molecular epigenetic determinants via their association with chromatin-related proteins^17^. Mouse BCNT gene, *cp27* was suggested to mediate early organogenesis and high level of its expression was demonstrated in developing mouse teeth, heart, lung and liver^18–20^. Thus, while there are sparse studies accessing the role of *cfdp1* in chromatin remodeling complex via the maintenance of chromosome organization^21,22^, little is known about not only its function and the mechanism it is involved in but also its role in heart physiology and morphogenesis. Recently, it was shown that the zebrafish *gazami* mutants carry a point mutation in the 3’ end of the gene, resulting in a truncated protein^23^. It is proposed that *cfdp1* controls neural differentiation and cell cycle in the cerebellum and retina, however its role in the heart was not studied.

Since, GWAS studies have revealed the implication of CFDP1 in the risk of CAD in humans, it is essential to unravel how *cfdp1* affects the proper development and function of the heart. For this purpose, we utilized zebrafish as model organism as it has emerged to be a valuable vertebrate tool in order to model human cardiovascular development and diseases^24–26^. The physiology of zebrafish development offers a precious advantage to study mutations that result in early embryonic lethality. For instance, mutations in the Cardiac troponin T (cTnT), also known as *sih* (*silent heart*) mutants exhibit a non-contractile heart phenotype, the *sih* embryos survive up to 5dpf (days post fertilization) as they uptake adequate oxygen through diffusion and are not dependent on a functional cardiovascular system and blood circulation until that developmental stage^27^. This ability allows the characterization of mutations that are embryonic lethal to other vertebrate models.

In this work, we aimed to study the previously unappreciated role of *cfdp1* during the development of the embryonic heart in order to elucidate its involvement in proper cardiac function. We showed evidence that cardiac expression of *cfdp1* is apparent during early developmental stages and plays an important role in myocardial trabeculation. As a consequence of *cfdp1* abrogation, embryos display heart dysfunction, contractility impairment and arrhythmias supporting its role on proper cardiac performance. In addition, mutant *cfdp1* embryonic hearts exhibit downregulation of Wnt signaling pathway in the mesenchymal cells of the inner valve region during valvulogenesis without affecting Notch activation in this process. Thus, loss of *cfdp1* affects directly or indirectly via cardiac function, valve development. *cfdp1*^−/−^ mutants and a percentage of heterozygous do not survive to adulthood as their heart develop severe arrhythmias and stop by 10 days post fertilization, suggesting a partially dominant phenotype of *cfdp1* loss of function.

## MATERIALS AND METHODS

### Fish housing and husbandry

Adult zebrafish were maintained and embryos were raised under standard laboratory conditions, at 28°C at a day-night cycle according to the Recommended Guidelines for Zebrafish Husbandry Conditions^28^. The zebrafish transgenic reporter lines used in the study were Tg(*myl7:GFP*) (also known as Tg(*cmlc2:GFP*)) for myocardium^29^, Tg(*fli1:EGFP*)^30^ for endothelial cells, Tg(*Tp1:mCherry*) for Notch-responsive cell^31^ and Tg(7x*TCF-Xla.Siam:nlsmCherry*)^32^ for Wnt-activated cells. CRISPR induced mutations to study cardiovascular genes, genotyping and adult handling of animals experimentation protocols were approved from the Bioethics and Animal committees of BRFAA and the Veterinary department of Attica region (numbers 247914 and 247916, 08/04/20) for facility EL 25 BIOexp 03. Embryos and larvae were anaesthetized by adding 0.4ml tricaine 0.4% (MS-222, Ethyl 3-aminobenzoate methanesulfonate salt) (Apollo Scientific, cat.# BIA1347) in 25ml E3 embryonic water at a final concentration of 0.0064% (v/v). Pigmentation of 24hpf embryos was prevented by adding phenylthiourea (PTU, Aldrich P7629) in E3 embryonic water at a final concentration of 0.003%

### gRNA and Cas9 mRNA synthesis

Identification and design of target sites to specifically knock-out *cfdp1* gene was performed by using the online CRISPR design tool CHOP-CHOP (https://chopchop.cbu.uib.no/). The selected target site is located in exon 3 and the sequence is 5’-CAGTAGGAGACATTGAAGAGCGG-3’. CRISPR gRNA mutagenesis was designed according to Jao et all., 2013 and briefly, the protocol used is as follows: The oligos were synthesized, annealed and cloned in pT7-gRNA (Addgene plasmid #46759). After E.coli transformation, selection of clones and identification of correct ones with diagnostic digestions to confirm the loss of BglII cutting site after successful insertion of target site, samples were Sanger sequenced. Following, the gRNA-vector was linearized by BamHI and putified. In vitro transcription of gRNA was performed using the T7 High Yield RNA Synthesis Kit (New England Biolabs, E2040S) and generated gRNA was purified. For making nls-zCas9-nls mRNA, the DNA vector pT3TS-nCas9n (Addgene plasmid #46757) was linearized by XbaI digestion and purified. In vitro transcription of capped Cas9 mRNA was performed using mMESSAGE mMACHINE T3 Transcription Kit (Invitrogen, AM13480).

### Microinjection in zebrafish embryos

Microinjections were performed either directly into one-cell-stage zebrafish embryos or in the yolk underneath the one-cell-stage embryo. The final concentration of injection mixture for the generation of *cfdp1* mutant line was: 300ng/μl Cas9 mRNA, 50-100ng/μl gRNA, 10% (v/v) Phenol Red, 20mM HEPES and 120mM KCL. *cfdp1* ATG-blocking morpholino was synthesized by GeneTools, LLC and the sequenced that was used is TCTGAATAATTCATTCTTGTGTCGT. The final concentration of antisense *cfdp1* morpholino used 0.4mM MO and 10%(v/v) Phenol red.

### Tail amputations

Adult zebrafish were immersed in fish system water with anesthesia (MS-222) for approximately 3 min. Then, caudal fins were amputated using fine scissors and zebrafish recovered by placing them in recovery tank and flushing the gills with fresh water. Caudal fins were placed in appropriate tube for further genotyping analysis.

### Sequencing and electrophoresis-based genotyping

DNA from zebrafish embryos and adult fish was extracted and target region of *cfdp1* was PCR amplified using flanking genomic primers: forward, 5- GGAGGCCTCAAACTGGTGGAG-3’ and reverse, 5-CTTCTGAGAGCTTGCACTTGG-3’. Amplicons were then prepared for Sanger sequence after product cleaning using ExoSAP (New England Biolabs, #M0293S, #M0371S). Alternatively, amplicons were visualized on a 2% agarose gel after diagnostic digestion to confirm the loss of SapI-cutting site inside the target site.

### RNA isolation and cDNA synthesis

Larvae were collected at different developmental stages, euthanized, transferred in 2ml tube containing 300μl TRI Reagent (Sigma-Aldrich, T9424) and homogenized. Extracted total RNA was reverse transcribed into cDNA using PrimeScript RT reagent Kit (TaKaRa RR037a) according to manufacturer’s instructions, using 500ng RNA per cDNA synthesis reaction.

### RT-PCR

For reverse-transcription PCR, synthesized cDNA was template for PCR amplification and primers that were used at this study are listed: *cfdp1*, forward: 5’- GAGACATTGAAGAGCGGCAG- 3’, reverse: 5’- CGACTTCTCCAGAGTGCTCA-3’; *actin*-2*b*, forward: 5’-CGAGCTGTCTTCCCATCCA-3’, reverse: 5’-TCACCAACGTAGCTGTCTTTCTG-3’. Quantification was performed in ImageJ Software and relative expression was normalized to *actin-2b* as a reference gene.

### Whole-mount *in situ* hybridization

Whole-mount RNA *in situ* hybridization (ISH) using *cfdp1* antisense probe was performed in embryos, according to The Zebrafish Book^33^. Primers for the generation of *cfdp1* probes were forward, 5’- GAGACATTGAAGAGCGGCAG-3’ and reverse, 5’- CGACTTCTCCAGAGTGCTCA-3’.

### Whole-mount immunohistochemistry

Zebrafish embryos were fixed with 4% paraformaldehyde overnight at 4°C and washed 3 times for 30min with PBS. Then samples were washed 3 times for 15 min with PBT (0.8% Triton X-100 in PBS) and incubated overnight at 4°C in phalloidin-633 (1:300 in PBT) for filamentous actin staining. Samples were then rinsed 3 times and washed 4 times for 15 min with PBT before mounting.

### Imaging

Zebrafish embryos were anaesthetized with 0.006% (v/v) Tricaine, placed dorsally on separate cavities of a glass slide (Marienfeld Superior, 10622434), mounted on 1,2% low-melting agarose and a drop of E3 embryonic water was added on top of semi-solidified mounting medium for maintenance of humidity. For *in vivo* imaging, fluorescent and brightfield videos of 10sec were recorded by microscope inverted Leica DMIRE2 with a mounted Hamamatsu ORCA-Flash4.0 camera. Confocal imaging was performed using a Leica TCS SP5 II on a DM 600 CFS Upright Microscope. The images were captured with the LAS AF software, analyzed in ImageJ Software and presented as maximum projection of a set of z-stacks for each stained tissue section.

### Adult zebrafish heart isolation and Histology

Adult zebrafish were euthanized in 0.016% tricaine containing 0.1M potassium chloride to arrest the heart chambers in diastole^34^. Images of whole hearts were captured using DFK2BUC03 camera from The Imaging Source mounted on SMZ1000 stereoscope. Then, adult hearts were fixed in 4% paraformaldehyde at 4°C overnight, washed three times with PBS, dehydrated in EthOH, and embedded in paraffin. Paraffin sections of 5μm thickness were performed using Leica RM2265 microtome. Haematoxylin and Eosin staining according to standard laboratory protocols. Images of stained sections were capture with Leica DFC500 camera mounted on Leica DMLS2 microscope.

### Estimation of cardiac function

High-speed videos of 30 frames per seconds and of 10sec duration taken under Leica DMIRE2 microscope were used to measure and calculate heart features. Heart rate (bpm, beats per minute) was calculated by counting the number of heart beats over the period of video and calculating the rate over 60sec. Derived from the still images of the videos, long-axis (lax) and short-axis (sax) of the ventricle during end diastolic and end systolic were measured and the average of three end diastolic and three end systolic per embryo was used to calculate ventricular volumes. Assuming that shape of ventricle is a prolate spheroid, the EDV and ESV were calculated using the following standard formula: V= (1/6) × π × (sax)^2^ × (lax) ^35^. Stroke volume was calculated by: SV= EDV-ESV. Cardiac Output was calculated by: CO= Heart rate × SV. Ejection Fraction was calculated by: EF (%) = SV/EDV × 100. And Shortening Fraction was calculated by: SF= (lax_(d)_-lax_(s)_)/lax_(d)_.

### Statistical analysis

Statistical differences between mutants and wildtype siblings were determined using two-tailed Student’s test. Statistical analysis and plotting were carried out in GraphPad Prism (version 5.03 for Windows). All data presented as mean ± SEM and *p*-value was considered significant *p ≤ 0.05, ** p ≤ 0.01, *** p ≤ 0.001.

## RESULTS

### 1. Expression profile of zebrafish cfdp1 gene during embryonic development

Albeit, the BCNT (Bucentaur) protein superfamily is highly conserved between species, their functional role remains unclear^10^. Previous studies, on model systems such as yeast *Saccharomyces cerevisiae* (SWC5)^13,36^, *Drosophila melanogaster* (YETI)^15^ and human cell lines (CFDP1)^22^ homologues have shown an important role during development by providing activity in chromatin remodeling organization. Two recent studies on zebrafish *cfdp1* linked the function of the gene with proper osteogenesis and craniofacial development focusing on the abundant *cfdp1* expression at the region of head^37^. Itoh et al., 2021, showed defective neuronal differentiation, particularly Vglut1 and Neurod1. In our study, we first analyzed the spatiotemporal expression of *cfdp1* during the development of zebrafish embryos, focusing on the cardiac area.

We performed whole mount *in situ* hybridization (ISH) with a specific *cfdp1* antisense RNA probe in wild-type embryos to investigate its expression pattern during development. At the first developmental stages, *cfdp1* expression is observed at the anterior part of the organism and at 120hpf the *cfdp1* expression is mainly detected at the cephalic region and the developing heart (Figure 1A). Control embryos that were hybridized with sense *cfdp1* RNA probe showed no staining pattern (data not shown). In addition, semi-quantitative reverse transcription PCR with total RNA extracted from wild-type zebrafish embryos at different developmental stages (5hpf – 120hpf) showed that *cfdp1* transcripts are detected from the first stages of development suggesting that maternal *cfdp1* mRNA is provided (Figure 1B). To further investigate the cardiac *cfdp1* expression at this later developmental stage, 120hpf ISH-stained embryos were collected, embedded in paraffin and cut in 5μm tissue sections via microtome. The histology analysis showed that *cfdp1* is expressed at the surrounding layer of the heart (Figure 1C). These data reveal, for first time, that *cfdp1* might play an important role in zebrafish developing heart.

**Figure 1:**
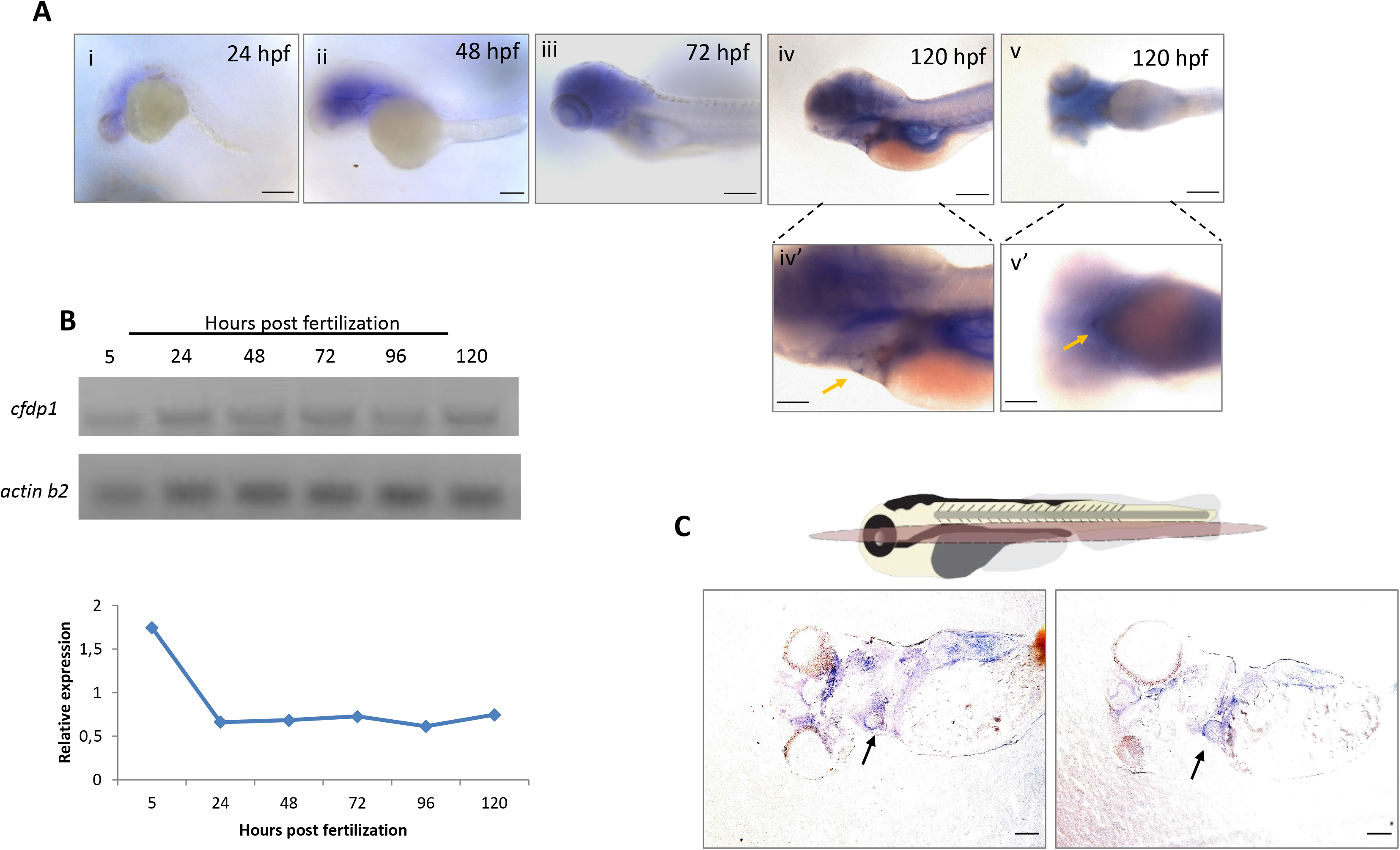
Expression analysis of *cfdp1* in different zebrafish development stages show that *cfdp1* is apparent during early development. A, Whole mount *in situ* hybridization of *cfdp1* in wild-type zebrafish embryos at different development stages (i-v). Higher magnification of iv and v shown in iv’ and v’ panels, respectively. Arrows point at the region of the heart. The expression of the gene is apparent from the 24 hpf and is restricted at the region of the head and the heart by 120 hpf. Scale bar (i-v) 150 μm, Scale bar (iv’-v’) 200 μm. B, Temporal expression analysis of *cfdp1* via RT-PCR compared to *actin b* house keeping gene. C, Upper: Illustration of frontal cutting plane of zebrafish, Lower: Paraffin sections of 120hpf ISH-stained embryos with *cfdp1* antisense RNA probe and *cfdp1* sense RNA (negative control). Arrows point at the stained embryonic heart. Scale bar: 50 μm

### 2. Silencing of cfdp1 expression reduces activation of Wnt pathway in the embryonic heart but Notch signaling remains unaffected

After the demonstration that *cfdp1* is also expressed in the zebrafish heart, it is important to address questions which concern the function of the gene during development. To achieve this, we first aimed to investigate *in vivo* the phenotypic characterization of embryos upon silencing of *cfdp1* expression via antisense morpholino oligonucleotide (MO)-mediated knockdown experiments. Therefore, we initially injected *cfdp1* translation-blocking MO in one-cell-stage wild-type embryos and then incubate them at 28°C up to 120hpf for monitoring. Figure 2A,A’ shows the phenotypic scoring of *cfdp1* morphants compared to the control sibling embryos. The majority of injected embryos develop pericardial oedema, from severe heart balloon shape to moderate oedema, along with craniofacial malformations, defects in otoliths and body curvature. The heart malformations of the *cfdp1* morphants phenotype affirmed the role of *cfdp1* in proper cardiac development and function.

**Figure 2:**
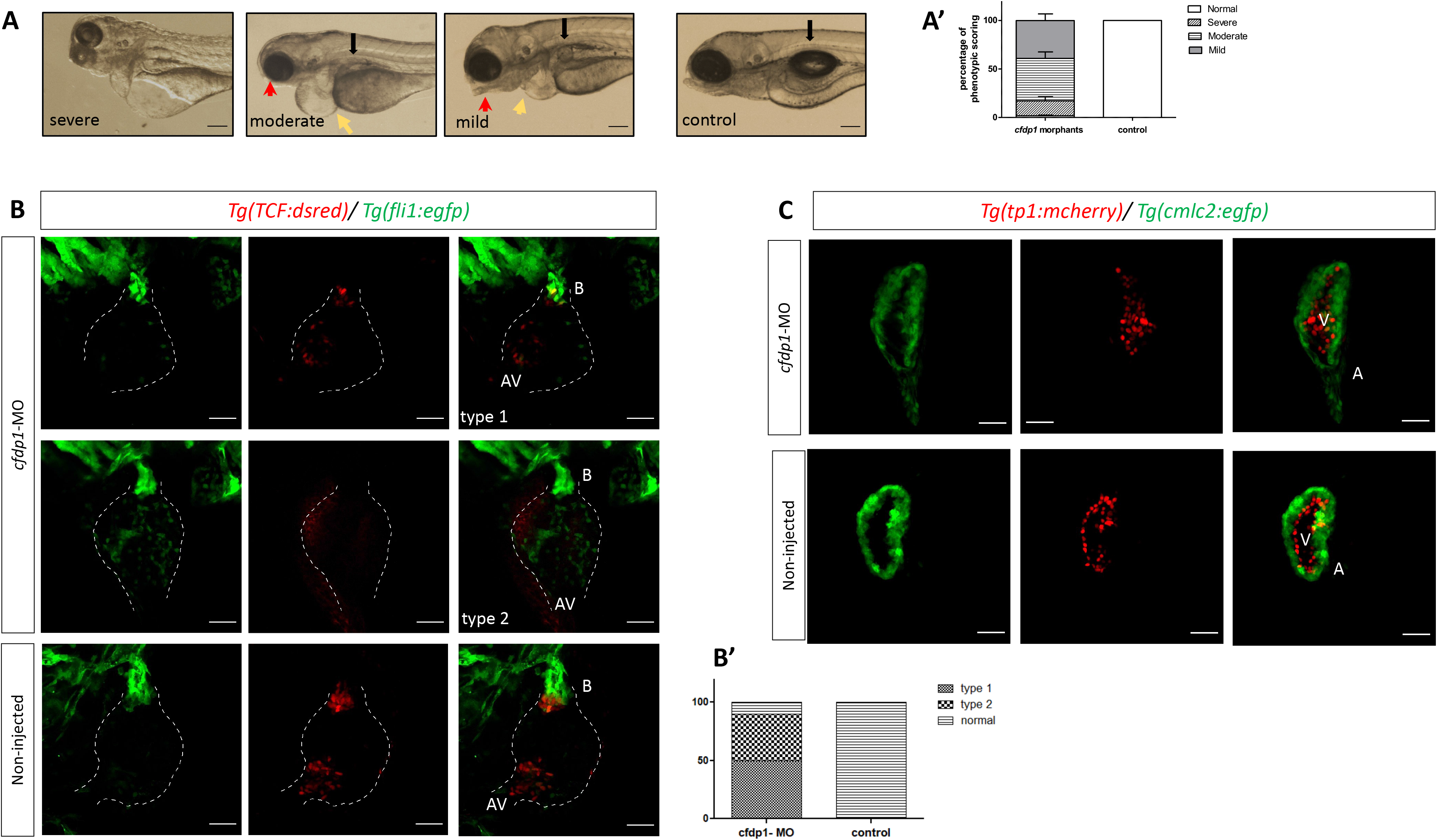
Silencing of *cfdp1* expression via morpholino microinjections. A, Stereoscopic images of representative 120hpf *cfdp1*-MO injected and uninjected control embryos. Black arrows point swim bladder, yellow arrows point pericardiac oedema, red arrows point mouth opening position. Scale bar 150 μm. A’, Quantification of phenotypic scoring via GraphPad Prism. B, Wnt/β-catenin activity is diminished in *cfdp1* morphants compared to the uninjected sibling controls. Max projection of z-stack confocal images of 72hpf *cfdp1*-MO embryos. Endothelial cells are labeled with green (Tg(*fli1:EGFP*)) and Wnt-activated cells are labeled with red (Tg(7x*TCF-Xla.Siam:nlsmCherry*)). B’ percentage of phenotypic scoring. AV, atrioventricular valve. B, bulbus arteriosus. Scale bar 150 μm. C, Notch signaling remains unaffected in *cfdp1* morphants compared to the uninjected sibling controls. Max projection of z-stack confocal images of 72hpf *cfdp1*-MO embryos. Ventricular cardiomyocytes are labeled with green (Tg(*myl7:GFP*)) and Notch-activated cells are labeled with red (Tg(*Tp1:mCherry*)). Scale bar 150 μm

Next, based on the importance of Notch and Wnt signaling pathways on the proper development, morphogenesis and function of the embryonic heart, we investigated whether these major regulator pathways are affected in *cfdp1* morphant embryos. We first investigated the Wnt/β-catenin signaling activity using the Wnt reporter line *Tg(7xTCF-Xla.Siam:nlsmCherry)*. It has previously shown that Notch and Wnt has different activity patterns since Wnt activity is primary located at the abluminal cells of the valves possibly mediating Epithelial-to-Mesenchymal Transition (EMT) of endocardial cells by increasing cell invasion during valve formation^39^. We crossed Tg(*fli1:EGFP*) (enhanced *GFP* expression under the endothelial specific promoter *fli1*) with Tg(7x*TCF-Xla.Siam:nlsmCherry*) and injected the *cfdp1* morpholino. We observed a significant downregulation of Wnt activity in the *cfdp1* morphants as they appear to have from reduced to complete absent signal of the Wnt reporter in the region of the heart (Figure 2B), (whereas reporter signal remains unaffected at the rest of the embryo, Supplementary Figure 1).

We then utilized the transgenic lines: Tg(*myl7:GFP*) to visualize myocardial cells and Tg(*Tp1:mCherry*) which indicate Notch-activated cells as the expression of *mCherry* is driven by the Notch-responsive element *Tp1*. *notch1b* is initially expressed throughout the endocardium of the heart and then becomes restricted at the valve-forming region and more specifically at the luminal endocardial cells of immature valve leaflets^38,39^. *cfdp1*-MO injected embryos showed no differences in the Notch reporter activation compared to control siblings at 72hpf (Figure 2C). Collectively, these findings show that *cfdp1* silencing affects Wnt/β-catenin but not Notch signaling indicating that *cfdp1* plays different role in these two stages and types of valvular cells.

### 3. Generation of zebrafish *cfdp1* mutant line

After the *in vivo* characterization of *cfdp1* morphants that suggested a role of *cfdp1* in the developing heart of zebrafish embryos which was previously unknown, we generated a knock-out *cfdp1* mutant line to circumvent any phenotypic discrepancies between morphants and mutants, previously reported^40^. Zebrafish *cfdp1* gene is located on chromosome 18 and consists of 7 exons, which encode for a protein of 312 amino acids (aa). In order to generate a mutation within the gene, we designed *cfdp1* guide RNAs (gRNAs) for CRISPR-Cas9-induced mutagenesis to target specific location according to published instructions (Jao et al...., 2013)^41^. We utilized the online tool CHOPCHOP (https://chopchop.cbu.uib.no/) and we targeted exon three as it scored at the highest ranking and efficiency rate (Figure 3A). The mixture of *cfdp1* gRNA and Cas9 mRNA was then injected at one-cell-stage embryos of the *Tg(myl7:EGFP)* line and the efficiency of the induced mutation was verified by Sanger sequencing of the flanking region around the target site of the injected embryos. Specifically, at 24hpf a pool of injected embryos was collected, DNA was extracted and a 350bp fragment across the target site was PCR amplified. The efficacy of guide RNA (and therefore the induction efficiency of somatic mutation) was assessed via DNA sequencing. Following, F0 fish were raised until adulthood when they were crossed with wild-type individuals to identify mutant founders and confirm that the induced mutation was transmitted to the F1 (Figure 3B).

**Figure 3:**
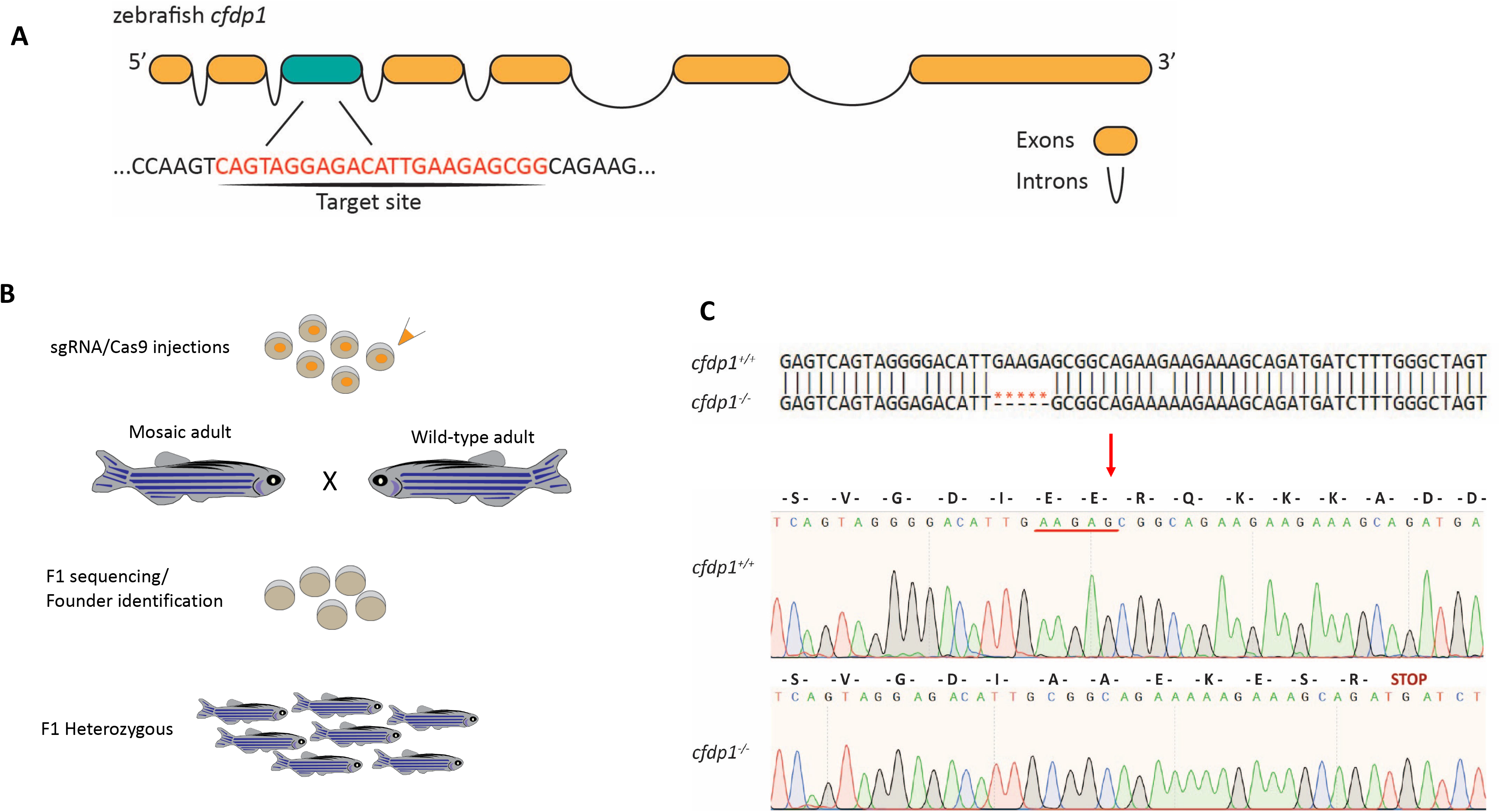
Generation of *cfdp1* zebrafish mutant line. A, Schematic representation of zebrafish *cfdp1* gene. For generation of CRISPR/Cas9-mediated mutant line, a target site in exon 3 was selected. B, Schematic representation of strategy for CRISPR/Cas9-mediated zebrafish line. The injection mix of gRNA /Cas9 in injected at the one-cell stage embryos. The crispants (F0 injected) grow until adulthood and are crossed with wild-type adults. The F1 generation is being genotyped in order to identify possible Founders of the line. After the identification, the corresponding F1 heterozygous generation in kept for further analysis. C, Upper: Nucleotide alignment between *cfdp1* mutant and *cfdp1* wild-type sequence. A 5bp deletion in *cfdp1* mutant is detected. Lower: Chromatogram of sanger sequencing of *cfdp1* mutant and *cfdp1* wild-type sequence and the corresponding aa they encode. In *cfdp1* mutant, at the point of DNA break, there is an insertion of seven novels aa and an early stop codon

A family of F1 carriers was selected, and the responsible mutated allele was characterized to be a deletion of five nucleotides. This caused a frameshift leading to an introduction of premature stop codon and as a result to the production of a truncated protein. More specifically, the deletion of AAGA before the PAM sequence of target site is predicted to result in a truncated product of 114aa, that have 107aa of the wild-type *cfdp1* protein and seven new aa before harboring the premature stop codon (Figure 3C). The highly conserved BCNT domain that resides at the C-terminal domain (exon six and exon seven) is also absent in the resulting truncated product. Additionally, the deletion of mutant allele led to the loss of the unique *SapI* cutting site in the surrounding area of the target site, which was subsequently used as a verification of *cfdp1* genotyping after diagnostic digestion of extracted DNA samples of *cfdp1* siblings (Supplementary Figure 2).

### 4. Zebrafish *cfdp1* mutants show impaired cardiac performance

Following the generation of *cfdp1* mutant line, we examined the phenotype of homozygous embryos and tried to raise homozygous adults. Intriguingly, homozygous mutant larvae did not reach adulthood and survived up to 10-15dpf. From a cfdp1^−/+^ cross we could only genotype adult heterozygous and wild-type. Thus, we proceeded to study the *cfdp1* role during development and early heart morphogenesis. Embryos from a *cfdp1*^*−/+*^ cross did not exhibit any gross phenotypic cardiac morphogenic malformations, but when monitored at 5dpf, we detected that a 27,8% of them suffered from cardiac arrhythmias (N=3, n=141) (Fig. 4A’). We, then carefully selected the individuals that have manifested the observed heart dysfunction and sequenced them, in order to identify their genotype. Interestingly, we discovered that not only homozygous *cfdp1*^*−/−*^ but also portion of heterozygous *cfdp1*^*−/+*^ siblings developed the observed heart dysfunction, while homozygous *cfdp1*^*+/+*^ (wild-type siblings) and also heterozygous *cfdp1*^*−/+*^ genotyped group were corresponding to the embryos without cardiac abnormalities. These findings highlighted the importance of *cfdp1* for proper heart function since homozygous mutants are larvae lethal, but also elude to a partially penetrant haploinsufficiency. In order to distinguish the homozygous *cfdp1*^*−/−*^ from the heterozygous individuals that are mimicking the severe phenotype of mutants, we sequenced retrospectively the sibling embryos (Fin Clipping) and proceeded further with the analysis (anterior embryo) based on the genotyping.

**Figure 4:**
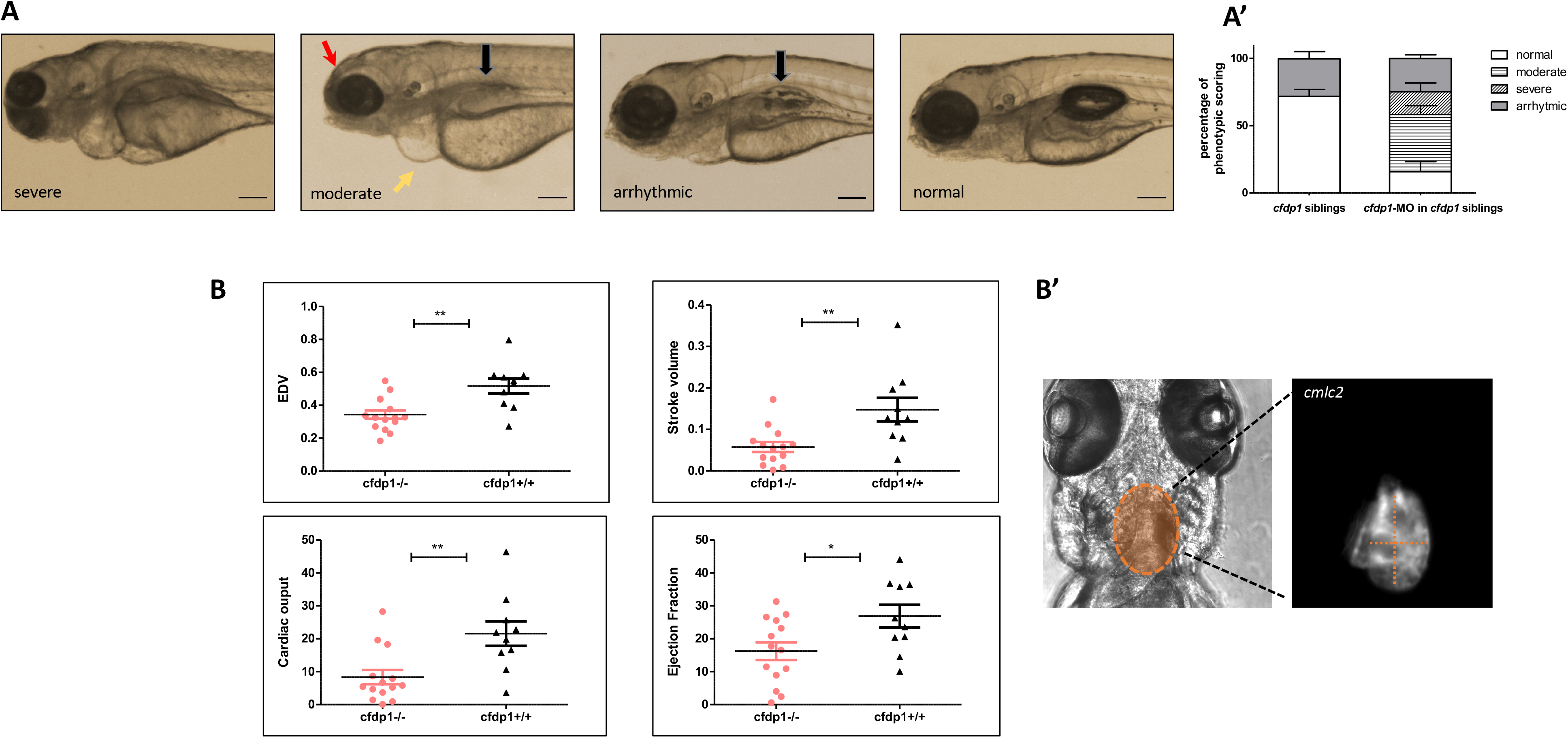
Impaired cardiac performance of *cfdp1* mutant embryonic hearts. A, *cfdp1*-MO injections in siblings embryos derived from cross between heterozygous *cfdp1* adult fish. Black arrows point swim bladder, yellow arrows point pericardiac oedema, red arrows point mouth opening position. A’, Quantification of phenotype scoring of *cfdp1* siblings (pool of all three genotypes: *cfdp1*^*−/−*^, *cfdp1*^*+/−*^, *cfdp1*^*+/+*^) and *cfdp1*-MO injected *cfdp1* sibling embryos. Scale bar 150 μm. B, Defective cardiac performance of 120hpf *cfdp1*^*−/−*^ embryos compared to their siblings *cfdp1*^*+/+*^ based on ventricular measurements after recording their heart rate. B’, Bright field and fluorescent image of *cfdp1* mutant embryos utilizing their Tg(*myl7:EGFP*) (also referred as *cmlc2*) background. Dashed lines indicate long and short ventricular axis, respectively.

Following, in order to assess the nature of the mutated *cfdp1* allele, we performed *cfdp1*-MO injections in one-cell stage embryos from F2 *cfdp1*^*−/+*^ adult individuals incross. Data showed in figure 4A that 22,3% of injected embryos developed arrhythmic hearts, 42,1% exhibited moderated phenotype with pericardial oedema, reduced size of head/eyes, malformations of mouth opening and flat or non-fully inflated swim bladder and 7,2% were scored as severe phenotype with gross abnormalities (N=4, n= 152). Therefore, the percentage of *cfdp1*-MO injected *cfdp1* sibling embryos that develop arrhythmic hearts is slightly reduced compared to the corresponding percentage observed to *cfdp1* siblings, while the appearance of moderate and severe phenotype scoring in *cfdp1*-MO injected *cfdp1* siblings is in accordance with the corresponding observed phenotypes in *cfdp1*-MO injected wild-type embryos (Figure 4A,A’).

We monitored *cfdp1*^*−/−*^ embryos and larvae and quantified several heart features through high-speed video imaging of single individuals (a method that was previously described by Hoage et al.,2012)^35^. Since, among the sibling group of embryos developing cardiac arrhythmias, homozygous *cfdp1*^*−/−*^ and heterozygous *cfdp1*^*−/+*^ were phenotypically inconspicuous, we recorded videos of all F3 *cfdp1* siblings at 120hpf acquiring brightfield and fluorescent images due to the fact that *cfdp1* mutant line was generated utilizing *Tg(myl7:EGFP)* reporter line and carries *myl7*-driven GFP expression for cardiomyocytes visualization. Embryos were then sacrificed and retrospectively analyzed after identification of genotype via DNA sequencing of single larvae. Remarkably, *cfdp1*^*−/−*^ showed significantly reduced end-diastolic volume and stroke volume as well as cardiac output and ejection fraction, compared to wild-type *cfdp1*^*+/+*^ siblings (Figure 4B). This strongly demonstrates that *cfdp1* abrogation inhibits proper ventricular function and stands as a strong effector in embryonic cardiac physiology.

### 5. Zebrafish *cfdp1* heterozygous develop variation in phenotype at embryonic stage

Due to the phenotypic heterogeneity in the group of heterozygous *cfdp1*^*−/+*^, we initially examined whether this is a result of considerable different levels of *cfdp1* expression within the corresponding genotype group. For this, we performed whole mount ISH with *cfdp1* RNA probe in a group of F3 *cfdp1* siblings 120hpf which contained a mixed of genotypes. Following, images of all ISH-stained embryos were captured and the expression levels of *cfdp1* were quantified via measuring ISH-staining pixel intensity that was analyzed using Fiji software^42^. This method represents an unbiased way of quantification of expression without prior knowledge of genotype. Therefore, after imaging, embryos were labelled and their genomic DNA was extracted and sequenced in order to retrospectively correlate their genotype to the quantified *cfdp1* expression levels. As expected, the heterozygous *cfdp1*^*−/+*^ showed a variation range of *cfdp1* expression between low and middle levels of intensity, which could explain the demonstration of the corresponding phenotypic variation within this genotype group (Figure 5A).

**Figure 5:**
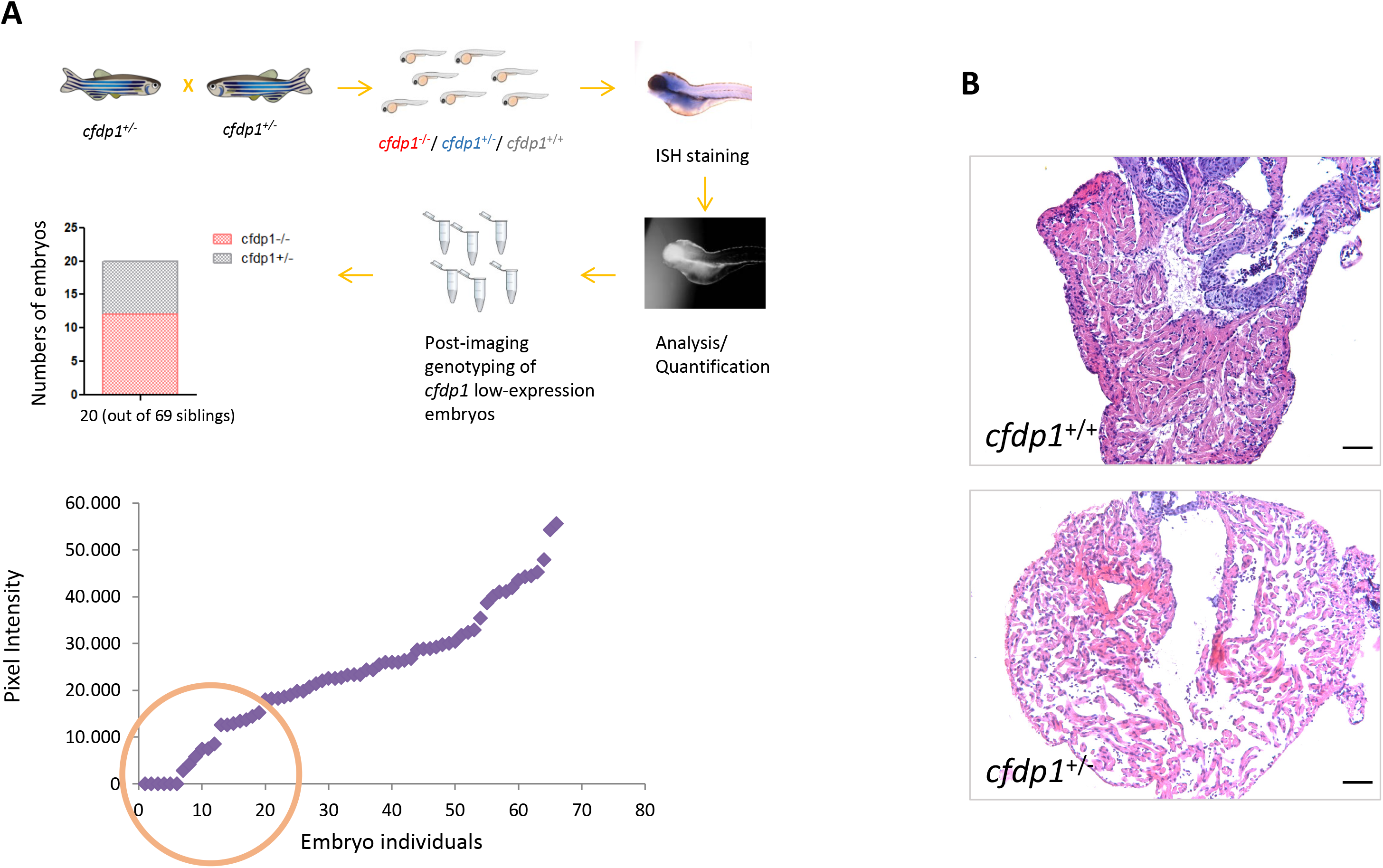
Study of *cfdp1*^*+/−*^ embryonic and adult hearts. A, Expression of *cfdp1* in *cfdp1* siblings (pool of three genotypes: *cfdp1*^*−/−*^, *cfdp1*^*+/−*^, *cfdp1*^*+/+*^). After performing *in situ hybridization* using *cfdp1* RNA probe in *cfdp1* siblings at 120hpf, ISH signal intensity was quantified. The *cfdp1* low- and no-ISH signal embryos were genotyped and it was confirmed that they corresponded to *cfdp1*^*−/−*^ and *cfdp1*^*+/−*^ embryos. B, H&E staining of 5μm paraffin embedded cardiac slices of *cfdp1*^*+/+*^ and *cfdp1*^*+/−*^ adult hearts. Scale bar 50μm.

Since, some heterozygous *cfdp1*^*−/+*^ reach adulthood, we then examine the structure of the heart of adult *cfdp1*^*−/+*^ compared to matched age *cfdp1*^*+/+*^ individuals. Here, adult fish were sacrificed and hearts were removed, sectioned and stained with Hematoxylin and Eosin for nuclear and ECM/cytoplasm staining, respectively. For accurate assessment, images of all heart sections were captured and analysis was performed on sections revealing the cardiac valves and the largest ventricular area. We observed that heterozygous *cfdp1*^*Δ/+*^ showed dilated ventricle, with thinner compact myocardium and sparse trabecular myocardium, with respect to wild-type control adult fish (Figure 5B). Overall, these findings show that the respective phenotypic variability of heterozygous is reflected to the levels of *cfdp1* expression and even so, the individuals that reach adulthood develop defects in heart morphology.

### 6. Zebrafish *cfdp1* mutants appear to downregulate Wnt pathway but Notch signaling remains unaffected

We wanted to verify the morpholino experiments and we investigated if *cfdp1* mutant embryos have defective Wnt or Notch signaling. For this purpose, we crossed adult heterozygous *cfdp1*^*−/+*^ (generated in Tg(*myl7:EGFP*) reporter line) with Tg(7x*TCF-Xla.Siam:nlsmCherry*) individuals and then raised the double positive egfp^+^/mCherry^+^, *cfdp1*^*−/+*^ offspring (for cardiomyocytes and *TCF*-activated cells visualization, respectively). 120hpf siblings from in-cross of adult *cfdp1*^*−/+*^/Tg(*myl7:EGFP*)/Tg(7x*TCF-Xla.Siam:nlsmCherry*) were screened under fluorescent microscope and double positive egfp^+^/mCherry^+^ larvae were genotyped (DNA samples derived posterior part of the embryo) while the anterior part was further processed and imaged to characterize the phenotype. As illustrated in Figure 6A, maximum projection of z-stack imaging reveals that Wnt pathway is significantly downregulated in mutant *cfdp1*^*−/−*^ compared to their wild-type *cfdp1*^*+/+*^ siblings, which is appropriately in line with the observed Wnt disruption in *cfdp1* morphants.

**Figure 6:**
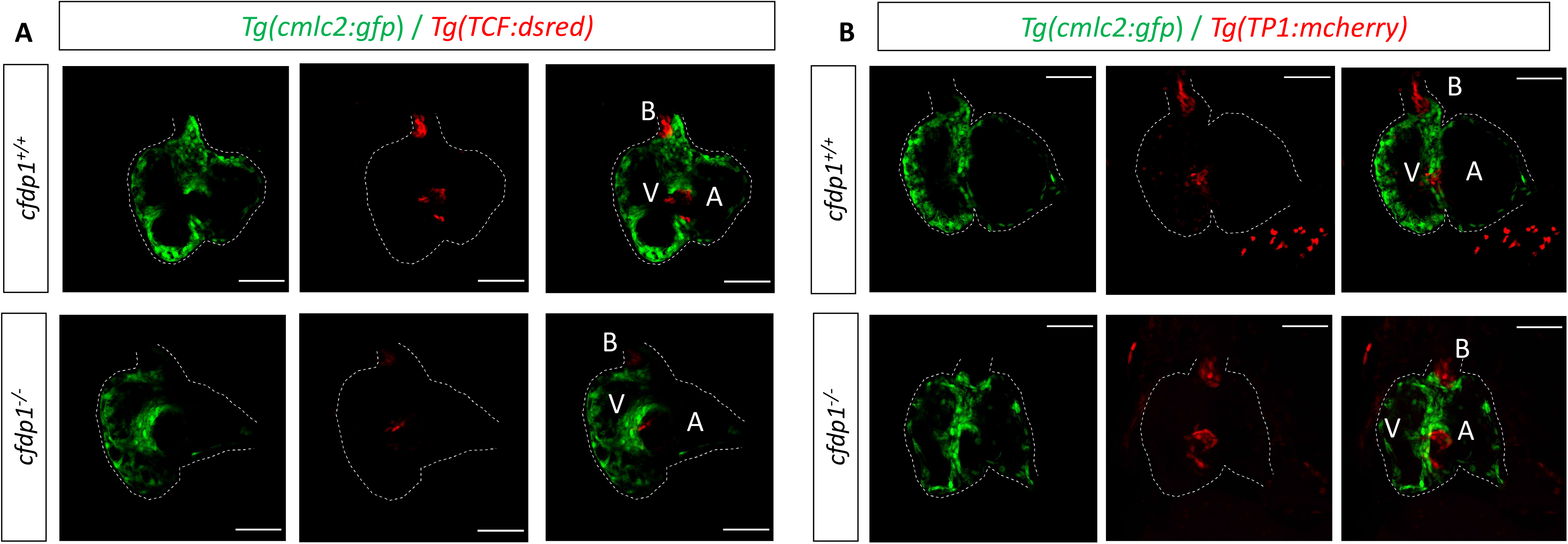
*cfdp1* abrogation show impaired Wnt/β-catenin signaling whereas Notch pathway remains unaffected. A, Confocal images of 120hpf *cfdp1* mutant and wild-type siblings expressing *Tg(Tcf:dsred)* in Wnt-activated cells and *Tg(cmlc2:eGFP)* in all cardiomyocytes. Scale bar: 50μm. B, Confocal images of 120hpf *cfdp1* mutant and wild-type siblings expressing *Tg(TP1:mcherry)* in Notch-activated cells and *Tg(cmlc2:eGFP)* in all cardiomyocytes. A: atrium, V: ventricle, B: bulbus arteriosus, Scale bar: 50μm.

We tested in a similar way the Notch signaling, which is also involved in valve formation with respect to the corresponding findings in *cfdp1* morphants. Adult heterozygous *cfdp1*^*−/+*^ with Tg(*Tp1:mCherry*) individuals were crossed with a *cfdp1*^*−/+*^ Tg(*myl7:EGFP*)/Tg(*Tp1:mCherry*) line. Remarkably, analogous to *cfdp1* morphants results, Notch activation pattern appears comparable to the wild-type *cfdp1*^*+/+*^ siblings (Figure 6B). In summary, Notch-expressing endocardial cells are differentiated while TCF-positive mesenchymal-like valvular cells exhibit lower activation levels.

### 7. Cardiac trabeculation in developing zebrafish ventricle is defective in *cfdp1* mutants

Prior studies have shown that orchestration of cardiac trabeculation is highly significant for the proper function of the heart and the survival of the embryo since defects during the complex morphogenic events occurring at trabeculation lead to embryonic lethality or adult dilated cardiomyopathies^43,44^. It has been also shown that zebrafish *erbb2* mutant embryos lack trabeculation but they develop normal valves^44^. To this end, we examined the levels of cardiac trabeculation in *cfdp1* mutant embryos at 120hpf when the entire length of luminal side of ventricle has developed extensive trabeculation. Single sibling *cfdp1* embryos were genotyped at 120hpf and groups of wild-type and mutant embryos were further stained with phalloidin for filamentous actin staining (Figure 7A). Interestingly, while *cfdp1* sibling wild-type embryos develop an extensive normal pattern of trabeculation, *cfdp1* mutant embryos exhibit less complex trabeculation (Figure 7B). This finding suggests the requirement of *cfdp1* for the proper initiation and formation of trabecular cardiomyocyte layer. As we have already shown, the *cfdp1* mutant hearts do not show signs of valve malformations, so the impairment of ventricle trabeculation is not a secondary effect to valvulogenesis defect. Taken together, our data show the *cfdp1* role specifically in cardiac trabeculation and cardiac function, while it is dispensable for valve formation.

**Figure 7:**
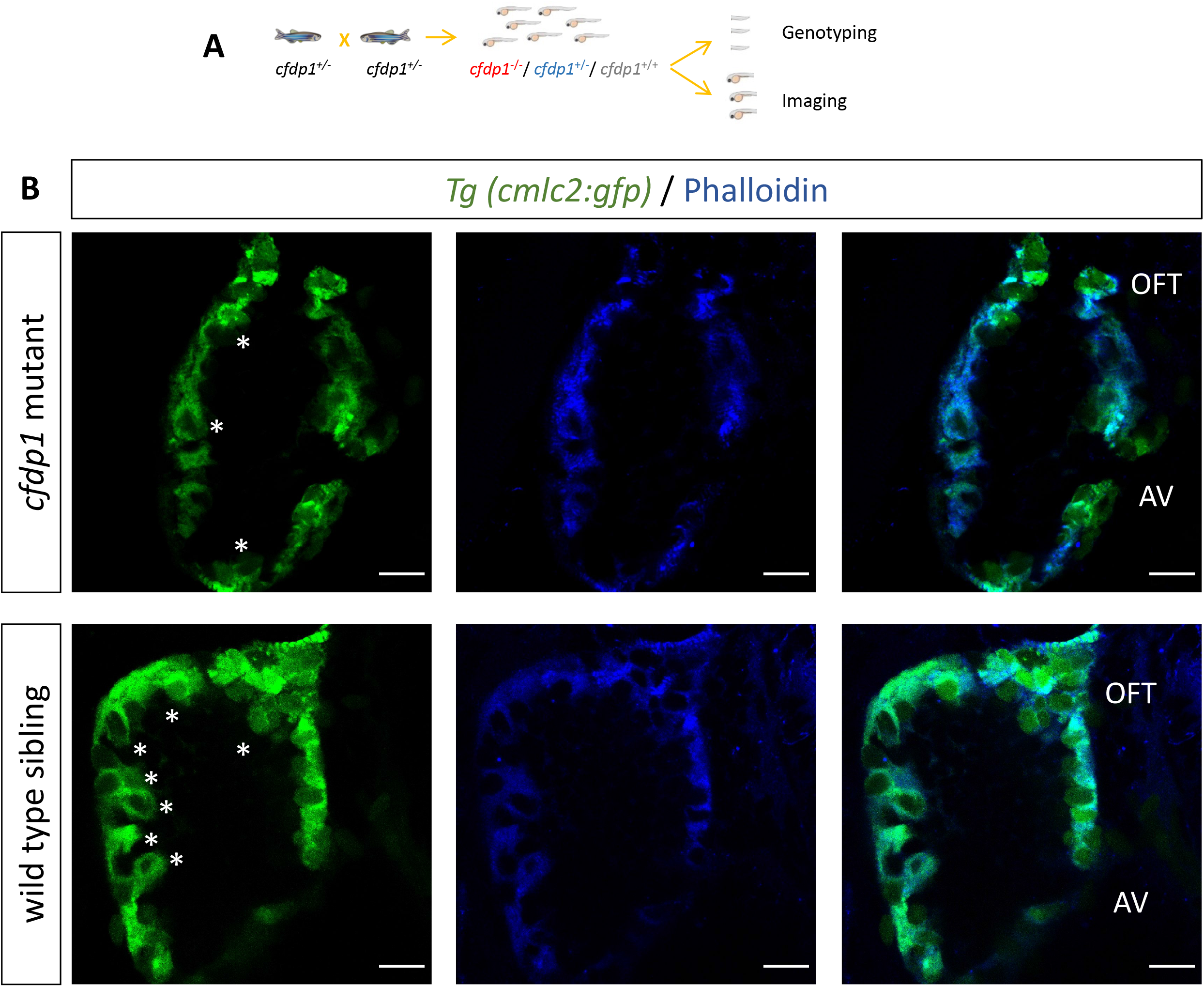
*cfdp1* is required for proper cardiac trabeculation A, Schematic representation of retrospective analysis. B, Single confocal plane of fluorescent phalloidin staining (actin filaments) in 120hpf *cfdp1* embryos, expressing *Tg(cmlc2:eGFP)* in all cardiomyocytes. Asterisks: trabeculae cardiomyocytes, AV: atrioventricular, OFT: outflow tract. Scale bar:50μm.

## DISCUSSION

In this study, we revealed for first time the essential role of *cfdp1* in cardiac development and proper function of the heart. We successfully generated a CRISPR/Cas9 - induced *cfdp1* mutant line by deleting five nucleotides around the PAM sequence resulting in alteration of reading frame, introduction of seven novel amino acids followed by an early stop codon and coding of a truncated protein product (missing the evolutionary conserved BCNT domain). This was achieved by targeting an oligonucleotide region on the third exon of zebrafish *cfdp1* orthologue and identification of mutant allele after sequencing. Our work demonstrated the cardiac dysfunction upon *cfdp1* abrogation which was reflected in decreased heart features of end-diastolic volume, stoke volume, cardiac output and ejection fraction. The *cfdp1*^*−/−*^ embryos do not reach adulthood as they die at approximately 10-16 dpf. We presented *in vivo* evidence of decreased ventricular trabeculation in *cfdp1* mutant hearts, while age-matched wild-type siblings showed normally developed trabecular network. In addition, *cfdp1* mutant embryos exhibited impaired contractility, bradycardia and arrhythmias, which is a characteristic observed in a small portion of *cfdp1* heterozygous embryos, as well. Interestingly, we showed that Wnt signaling in mesenchymal valvular cells is downregulated in *cfdp1* mutant hearts while they do not affect Notch activation in the atrioventricular boundary and the initiation of valve formation.

### Providing a valuable novel tool for phenotypic and functional characterization of *cfdp1* gene

Biochemical and functional analysis of CFDP1 (hBCNT/CFDP1) in human cell lines (HeLa, U2OS and MRC5) identified two isoforms of 50 kDa and 35 kDa (spliced variants) found in the nucleus of the cells^22^. The same study suggested that the 50 kDa variant has chromatin-binding activity (while the shorter isoform obtains different characteristics) and plays an important role in chromatin remodeling and organization affecting the progression of cell cycle. Interestingly, the truncated construct Flag-CFDP1-Nt containing only the N-terminal and lacking the conserved BCNT domain (C-terminal region) was able to enter the nuclei but lost the chromatin binding activity resulting in a defective truncated product^22^. In our zebrafish model, the mutated *cfdp1* allele generated via CRISRP/Cas9 system lacks also the BCNT domain since early stop codon is inserted close to the N-terminal region of the protein product. Based on the fact that this domain is highly evolutionary conserved between species, it is expected that z*cfdp1*^−/−^ loses the chromatin binding activity as well, and therefore its subcellular mechanical function but this needs to be further clarified and confirmed.

The characterization of *cfdp1* orthologous has been largely unexplored. Both *in vitro* and *in vivo* studies on yeast (*Saccharomyces cerevisiae*) BCNT orthologue SWC5, have shown that SWC5-deleted mutants lack SWC1-mediated Htz1 histone replacement suggesting that SWC5 is required for chromatin remodeling which can impair transcription and other cellular responses^36,45^. In the same context, studies in *Drosophila melanogaster* have demonstrated YETI, the BCNT member to be a multifaceted chromatin protein found in cell nuclei whereas its depletion in *Yeti* mutants leads to lethality before pupation^15^. *Yeti* binds to chromatin via its BCNT domain and interacts with both H2A.V variant and HP1a and it is proposed that YETI participates in the control of transcription initiation or the chromatin integrity. The first evidence of heart localization of BCNT genes during development comes from embryonic mouse studies revealing that CP27, the BCNT orthologue, is expressed in the developing heart (E8-E10), as well as organs like brain neuroepithelium, teeth, retina of the eye, otic vesicles, cerebellum and periosteum developing bones and in most cases, CP27 signal expression is found in epithelial-mesenchymal boundary in developing tissues (except dental pulp and periosteum)^18^. Later on, CP27 loss of function in mouse embryonic fibroblast cell line BALB/c 3T3 showed reduction in fibronectin matrix composition and redistribution of extracellular matrix (ECM) organization, suggesting that CP27 has a regulative effect on ECM and cellular changes^20^. It is known that ECM synthesis and remodeling promotes trabecular rearrangements and trabecular network growth in non-compaction cardiomyopathy (NCC) mouse model and it has been shown that fibronectin exhibit similar pattern to *Has2*, *Vcan* and CD44 (a hyaluronan receptor), which are ECM synthesis genes essential for trabeculation^46^. Therefore, a possible involvement of CP27 in ECM remodeling during ventricular trabeculation in mice, should also be investigated.

Genome-wide association studies have unraveled multiple genome loci associated with human diseases. A recent study performed deep transcriptomic analysis of genotyped primary human coronary artery smooth muscle cells (HCASMCs) and coronary endothelial cells (HCAECs) from the same subjects and analyzed GWAS loci associated with vascular disease and CAD risk, in these two coronary cell types^47^. Researchers found *CFDP1* (along with *YAP1* and *STAT6*) for HCAECs that passed the 5% false discovery level (FDR) correction at the gene level which associates *CFDP1* with artery disease traits^47^. Another study which applied a 2-stage discovery and replication study design with more than 15000 individuals, identified an association of a novel SNP in the last 3’ intron of *CFDP1*, rs4888378, with carotid intima-media thickness (cIMT), an established marker for subclinical atherosclerotic cardiovascular disease^9^. A different study identified another *CFDP1* variant, rs3851738, as CAD-associated locus after analysis from UK Biobank and CARDIoGRAMplusC4D 1000 Genomes imputation study, and following ‘phenome-wide association study’ (PheWas) correlated this variant with systolic blood pressure^48^. In the same context, GWAS studies have shown correlation of human *CFDP1* with aortic root diameter, as well as CAD risk^6,49^.

Thus far, *in vivo* studies clarifying specifically the involvement of *cfdp1* in cardiac development are voided. A detailed phenotypical and functional analysis of the GWAS-derived *CFDP1* is essential to shed light to the way of action and its determinant role in cardiovascular physiology. The present work provides evidence for first time about the fundamental effect of *cfdp1* in proper heart morphogenesis and function in zebrafish. The observed phenotype of bradycardia and arrhythmias is an observation with potential clinical relevance for CFDP carriers and their risk to develop CAD.

### *cfdp1* knockdown and knockout zebrafish models demonstrated similar but not identical results

Targeted knockdown of genes via MO injections is distinguishable from stable genetic lines which inherit the induced change, since MOs are gradually degraded within few days and therefore result in a transient effect. Despite that fact, knockdown approach in zebrafish remain an *in vivo* phenotypic assay to investigate the effect of gene depletion upon blocking of their expression. Our data showed that *cfdp1* morphants develop phenotypic abnormalities, such as pericardial oedema, craniofacial malformations and hypoplastic swim bladder (arrest of swim bladder inflation has been proposed to be a secondary event to heart failure, since in *silent heart* morphants that lack heart contractility, heart-specific constitutively activated AHR signaling and TCDD-exposed zebrafish models which develop heart failure, the swim bladder development is inhibited in the same manner^50^). Interestingly, the *cfdp1* mutants exhibit more mild phenotypic characterization by developing arrhythmic embryonic hearts but not pericardial oedema or extreme craniofacial disorders compared to *cfdp1* morphants at 120hpf. At the same context, when we investigated the effect of *cfdp1* depletion in Wnt signaling pathway at *cfdp1* morphant hearts, we observed major reduction in signal intensity or even complete blockage of expression pattern in Wnt-activated cells of Tg(7x*TCF-Xla.Siam:nlsmCherry*) reporter line, while *cfdp1* mutant hearts show strong inhibitory effect without total silence of Wnt pathway. The differences in manifest of *cfdp1* depletion between knockout and knockdown embryos could possibly be accounted for by the activation of a genetic compensation response, which has been previously proposed to explain phenotypic discrepancies in morphants and mutant models^40^.

### Variation of *cfdp1* heterozygous phenotype manifest

The generation of stable *cfdp1* zebrafish mutant line resulted in the induction of a deleterious mutation caused by harboring a premature termination codon (PTC) in *cfdp1* sequence. Detailed phenotypic study of *cfdp1* sibling embryos unveiled the arrhythmic hearts of *cfdp1*^*−/−*^ embryos. Notably, the same phenotype emerged in a range of heterozygous *cfdp1*^*−/+*^ embryos that made them undistinguished from the *cfdp1* mutants. We further investigated whether this could be a result of variation in *cfdp1* expression levels and indeed, we detected differences in signal intensity within *cfdp1*^*−/+*^ embryo pool, suggesting that this could modulate the phenotypic variation of heterozygous zebrafish. Our data support the existence of heterogeneity (variation of phenotype) in heterozygous *cfdp1* siblings (same genotyping group) and the possible correlation of wild-type/mutated copies and phenotypic outcome. A proposed scenario for this variation holds on the activation of quality control nonsense-mediated mRNA decay (NMD) that targets flawed messenger RNAs. Since, our *cfdp1* mutation induces a PTC that is not at the last exon and is ~ 50 nucleotides upstream of the last exon-exon junction, it is well assumed that triggers NMD machinery^51^. It is generally known that, NMD is a surveillance pathway that degrades transcripts containing PTCs in order to maintain transcriptome homeostasis^52,53^. Although NMD plays a beneficial role by limiting the dominant-negative effect of mutant proteins, there is a variation in the efficiency of NMD activity in cell-, tissue- and transcript-specific differences that modulates the manifestation of a disorder^52,53^. Interestingly, it has also been suggested that NMD variation potentially leads to different clinical outcomes in individuals carrying the same PTC-containing mutated transcript^54^. For instance, patients containing the same mutation in *X*-chromosome develop markedly different phenotypes (Duchene Muscular Dystrophy and Becker Muscular Dystrophy, respectively) upon differentially activation of NMD, allowing the accumulation of truncated protein in one case^54,55^. Thus, the efficacy of NMD could vary between individuals and acts as potential modifier of disease phenotype. Therefore, the observed variability between *cfdp1*^*−/+*^ individuals could also be a consequence of incomplete NMD resulting in *cfdp1* haploinsufficiency and heterogeneity observed in heterozygous carriers, but it needs to be further investigated.

### The role of *cfdp1* in ventricular trabeculation and cardiac function

After cardiac chamber formation, cellular remodeling leads to a formation of an intricate architecture through the initiation and growth of ventricular trabeculation. Numerous of signaling pathways in endocardium, myocardium and cardiac ECM are involved in the regulation of this process, such as Notch^46,56^, Semaphorin 3E/PlexinD1^57^, angiopoietin/Tyrosine kinase with immunoglobulin-like and EGF-like domains (Tie2)^58^, Bone morphogenic protein (BMP)^59^, EphrinB2/EphB4^60^, and most importantly Neuregulin (nrg) signaling which operates through ErbB receptor tyrosine kinase. E10.5 days postcoitum (dpc) *nrg1*^*−/−*^ mice suffer from severe impaired trabeculation, as well as increased apoptotic levels at the region of the head, reflecting its role also in cranial neurogenesis^61^. Similarly, null mutations in ErbB2 and ErbB4 result in abrogation of ventricular trabeculation that lead to lethality between E10.5 and E11.5 in mice^62,63^. Zebrafish *erbb2* mutant embryos lack cardiac trabeculation and develop progressive cardiac dysfunction and fatal heart failure, showing the functionally conserved role of Nrg/ErbB signaling in heart morphogenesis^44^. Interestingly, *erbb2* mutants exhibit normal valve morphogenesis, indicating a direct and cell-autonomous regulation of ErbB2 in cardiac trabeculation. In addition, while in *nrg1* zebrafish mutant larvae trabeculation appears unaffected and *nrg1*^*−/−*^ survive to fertile adults, *nrg2a* (another member of Nrg family) mutants hearts fail to form trabeculation and suffer defects similar to *erbb2* mutants^43^. Notably, *nrg2a*^*−/−*^ are recognized morphologically by their aberrant jaw and swim bladder inflation disorders, reminiscing of the phenotypic characterization of *cfdp1* morphant embryos. In accordance to what it was observed in *erbb2* mutants, *nrg2a* zebrafish mutants develop normal atrioventricular (AV) valves, indicating that Nrg2a/ErbB2 is dispensable for AV valve formation and it is required for proper cardiac trabeculation. Interestingly, zebrafish *tomo*-seq genome-wide transcriptional profiling^64^ reveal similar expression pattern of *cfdp1* (previously also known as *rltpr*) and *nrg1* in regenerating heart 3 days after injury, indicating a possible functional association between the two genes. Whether c*fdp1* mechanism of action and its role in trabeculae cardiomyocytes regulation crosslinks with Nrg signaling pathway remains to be further investigated.

We have shown that *cfdp1* zebrafish mutants suffer from impaired trabecular network. Defects of this complex cardiac remodeling lead to embryonic lethality, which illustrates the importance of this process and the need to fully unravel the signaling molecules regulating the trabeculation in cardiac development. Mechanical forces and contractility are also important factors for the proper trabeculation network formation. Both reduction of blood flow in *weak atrium* (*myh6*)^65,66^ mutants and disrupted contractility in *silent heart* (*tnnt2a)*^67^ mutants result in severe defects in trabeculation, as well as *tnnt2a* morphants that do extend ventricular protrusions but they are less stable and frequently retract^68^. Disorder in trabeculae layer shown in *cfdp1* mutant embryos could be a secondary event of reduced contractility which is demonstrated by reduced stroke volume and ejection fraction cardiac performances. It would be interesting to utilize the zebrafish transgenic line *Tg(cmlc2:gCaMP)*^67^, a cardiac-specific fluorescent calcium indicator line to monitor the cardiac conduction signal travel in *cfdp1* mutants in order to further investigate the correlation of contractility and trabeculation in *cfdp1* embryonic mutant hearts.

Since previous studies have illustrated that Notch and canonical Wnt/β-catenin signaling pathways expressed in endocardial cell are influenced by blood flow and contractility^39^, we investigated how modulation of contractility in *cfdp1* mutants affects the activation of these major molecular pathways. We demonstrated that Wnt/β-catenin signaling reporter line exhibited disrupted expression pattern, while Notch-activated cells in the corresponding reporter line didn’t show any effect. The different activities of Notch and Wnt/β-catenin observed in *cfdp1* mutant hearts indicate the composition of two different cell subsets, in accordance to previously reported Notch-activated luminal AV cells and Wnt/β-catenin-activated abluminal AV cells during valve formation^39,69^. Moreover, it has been shown that although Notch and Wnt signaling intersect in order to promote the TCF-positive endocardial cells ingression into cardiac jelly during valvulogenesis, inhibition of Erk5-Klf2 pathway impairs canonical Wnt signaling without affecting Notch nor Dll4 activation in atrioventricular endocardial cells, confirming that these pathways are regulated independently^69^.

Cardiac conduction system is composed by pacemaker cells in sinoatrial junction, atrioventricular node and ventricular conduction system and canonical Wnt pathway has been implied to contribute during specific stages of conduction^70^. Canonical Wnt5b signaling has been reported to play an important role in heart contractility by promoting pacemakers cardiomyocytes differentiation transcription factors Isl1 and Tbx18 and inhibiting Nkx2.5, both in zebrafish and human pluripotent stem cells (hPSCs)^71^. Likewise, Wnt signaling activation (via Wnt3 ligand) promotes pacemaker lineage in mouse and human embryonic stem cells^72^. In *Isl1*-deficient zebrafish and mouse embryos, there is a progressive failure of contractility leading to arrhythmias and bradycardia^73^ and it is reported that canonical Wnt/β-catenin signaling in zebrafish is activated in *isl1*^*+*^ cells in sinoatrial region affecting the control of heart rate^74^. In addition, Wnt/β-catenin signaling in AV canal regulates specific electrophysiological properties of AVC and AV node by slowing down conduction velocity^75^. We reported that *cfdp1* embryonic mutant hearts exhibit arrhythmias, a phenotype indicating defects in contractility and pacemaker activity. Having highlighted the significant role of Wnt in regulating pacemaker development in zebrafish, *cfdp1* seems to function in regulatory mechanism upstream of Wnt pathway involved in cellular specification of conductivity. The mechanism of how *cfdp1* cooperates with canonical Wnt/β-catenin signaling remain to be elucidated.

In summary, the CRISPR/Cas9-induced *cfdp1* zebrafish mutant line provides an unprecedented tool to unveil novel mechanism of regulating cardiac physiology and function as well as ventricular trabeculation during embryonic development.

## Funding

This research is co-financed by Greece and the European Union (European Social Fund-ESF) through the Operational Programme «Human Resources Development, Education and Lifelong Learning» in the context of the project “Strengthening Human Resources Research Potential via Doctorate Research” (MIS-5000432), implemented by the State Scholarships Foundation (ΙΚΥ)

## Notes

### Competing Interest Statement

The authors have declared no competing interest.

