## Supplementary figures for "*Cfdp1* is Essential for Cardiac Development and Function"

SF1: *cfdp1* silencing via morpholino knockdown does not affect Wnt signaling-expression in posterior embryo.
SF2: Diagnostic test after digestion of DNA samples with *SapI* restriction enzyme. KO, knockout, Het, heterozygous, WT, wild-type.
